## Supplementary material for "Convergent Evolution and B-Cell Recirculation in Germinal Centers in a Human Lymph Node": Supplemantary Materials

### Supplementary Information for: Germinal Centers Convergent Evolution and B-cell Recirculation in Human Lymph Nodes

### 1 Mutations analysis

#### 1.1 Mutations matrices

For each sample, we first separate sequences into functional and non-functional groups. Then, we compute the mutation matrices for both groups, in which the transition between each nucleotide is quantified as

$$\mathcal{M}_{ACTG} = \begin{pmatrix} \text{NA} & a & b & c \\ d & \text{NA} & e & f \\ g & h & \text{NA} & i \\ j & k & l & \text{NA} \end{pmatrix}, \tag{1}$$

where each number $(a..l)$ represent the proportion of mutations from one nucleotide to another. The matrix is normalized such that $\sum_{k=a}^{l} k = 1$. Then, we define the distance between two matrices as the average absolute difference between each of their components:

$$dist_{ACTG}(\mathcal{M}_1, \mathcal{M}_2) = \frac{1}{12} \sum_{k=a}^{l} |k_1 - k_2| \tag{2}$$

Matrices and other parameters were obtained by submitting sequences into the AR-galaxy Web tool and using the SHM & CSR pipeline [1]. Computing the distance matrix for all considered samples reveals only a minimal difference across samples, for both the functional and non-functional sequence groups, with a distance always lower than 5 % across all groups (Figure S1A).

#### 1.2 Replacement to silent mutation ratio (R/S)

We compute the replacement to silent mutation ratio (R/S) in both CDR and FR regions in sequences of each germinal center by submitting sequences into the AR-galaxy Web tool and using the SHM & CSR pipeline [1]. As a high ratio indicates that replacement mutations are being selected over silent mutations, it is expected that the ratio positively correlate to the selection pressure undergoing in an GC environment. We compared the R/S ratio over three different groups, functional vs non-functional sequences (N vs NF), FR mutations vs CDR mutations (FR vs CDR) and sequences from the top 100 dominant clones vs singletons (D vs S) (Figure S1B), and computed their statistical significance with the T-test [2]. We have chosen top 100 clones in order to have enough sequences to do statistical analysis for the non functional clones. In order to avoid the skewing of the SHM spectrum due to the expansion of the dominant clones, we assigned one sequence per clone, defined as the most frequent one within each clone. Furthermore, we studied only the singletons that are found consistently in both replicates of each GC. Mutations in the CDR regions had a R/S ratio approximately twice as high as in the FR region except for non functional dominant clones (average R/S of 2.8 vs 5.5, $p < 0.004$), showing the expected major difference in selection pressure between FR and CDR regions. On the other hand, mutations in nonfunctional top 100 clones show non significant difference ($p = 0.62$) between FR and CDR R mutations, proving that the non-functional clones are not under selection pressure. Regarding sequences from functional dominant clones and singletons, the R/S ratio was observed to decrease in and CDR regions as less frequent clones were considered showing the different magnitude of selection pressure in different classes of dominance (Figure S1C). We have to take into account that approximately 25-30 clones were expanded in each GC ( $frequency > 1\%$) that observation can explain the

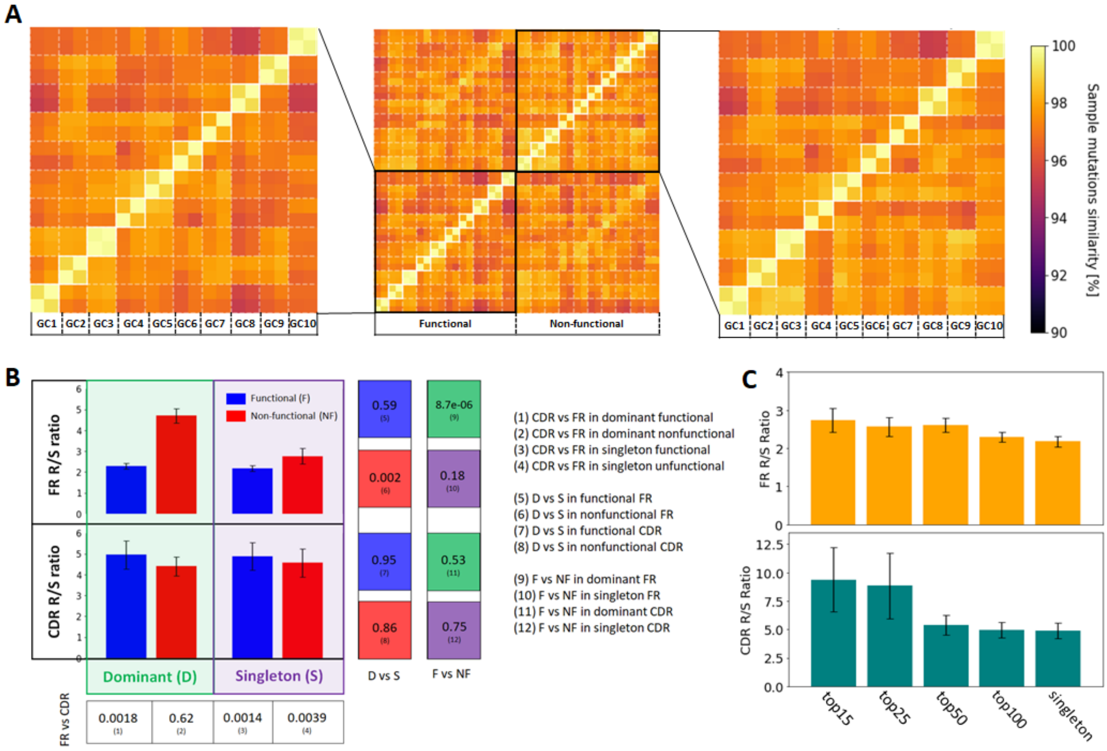

Figure S1: Mutations in functional and non-functional sequences across samples. (A) Sample distance matrix in term of mutation transition tables, where the similarity is defined as $1 - dist_{ACTG}$. (B) Replacement/silent ratio in CDRs and FR in functional and nonfunctional singletons and dominant clones, averaged over the 10 GCs. The error bar represent the standard error of the mean R/S estimation ($std/\sqrt{10}$). $p$-values (T-test) for each relevant comparisons are provided on the right side (1-12), where statistically significant comparisons ($p < 0.01$) are highlighted in green. (C) Average replacement/silent ratio in CDRs and FR in functional clones, selected as the top 15, 25, 50 and 100 most dominant clones in each GCs, respectively, as well as singletons.

fact that we do not observe significant difference in the CDR R/S ratio between the top 100 clones and the singletons (Figure S1B).

#### 2 Optimal threshold for clonal identification

During clonal identification, we separate sequences into distinct clonal groups when their respective junction nucleotide sequence identity is below a certain threshold. To determine the optimal cutting threshold, we combine the bimodal distance-to-nearest distribution model previously described by [3], and the negation sequences approach proposed by [4].

The first method relies on the fact that sequences belonging to one clone (non singletons) are similar to other sequences from the same clone, while sequences with no clonal relatives in the data set (singletons) will have a higher distance to the nearest junction in the dataset. The distance to nearest distribution will thus consist of two combined distributions, one from singletons and the other from non-singletons, that can be fitted with the sum of two gamma distribution. The optimal threshold would then be at the intersection of the two fitted distributions. Under the assumption that clones cannot span multiple individuals [4], the second approach samples sequences from multiple unrelated individuals (negation sequences), and uses them to estimate the specificity of the clonal identification framework for a given threshold.

We computed the normalized Hamming distance between all junctions pairs with the same length, where only a single replicate of each junction was considered. Then, the distribution of distances between the sampled sequences and their closest counterpart within the sample, as well as within the negation sequences, was computed. The negation table was built from randomly selected functional BCRs that express either IGHG or IGHM constant regions and do not have any indels, provided by [4].

As highlighted in Figure S2, the singleton and non-singleton distributions are clearly distinguishable, and the singletons distribution matches the negation distribution well. By setting our threshold at the intersection of the singleton and non-singleton distribution, we obtain a threshold of 0.16, which leads to a false positive rate (sequence pairs wrongly assigned to the same clone) of $\sim 1\,\%$ according to the fraction of negation sequences below the threshold.

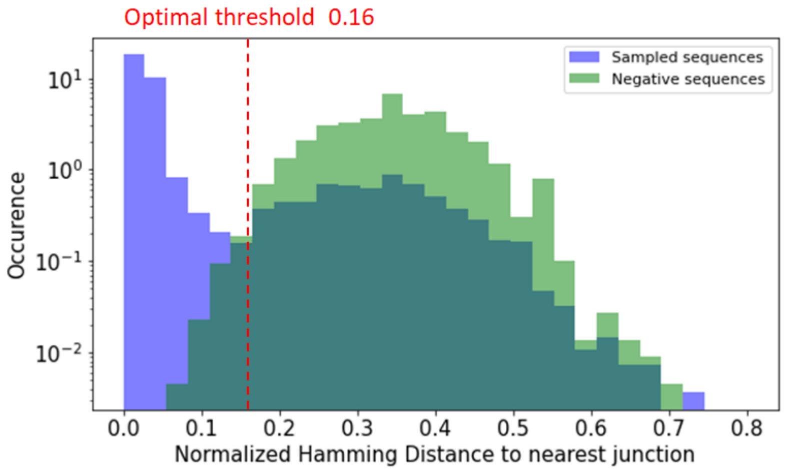

Figure S2: Distance to nearest distribution to both negation sequences (green), and sequences within the same sample (blue).

### 3 V and J gene analysis

To further illustrate and quantify the V and J gene selection process within the GC, we consider the VJ gene combination heatmap (Figure S3), defined as the V gene abundance histogram collected over multiple ranking thresholds of the definition of expanded clone (top10 to top 400).

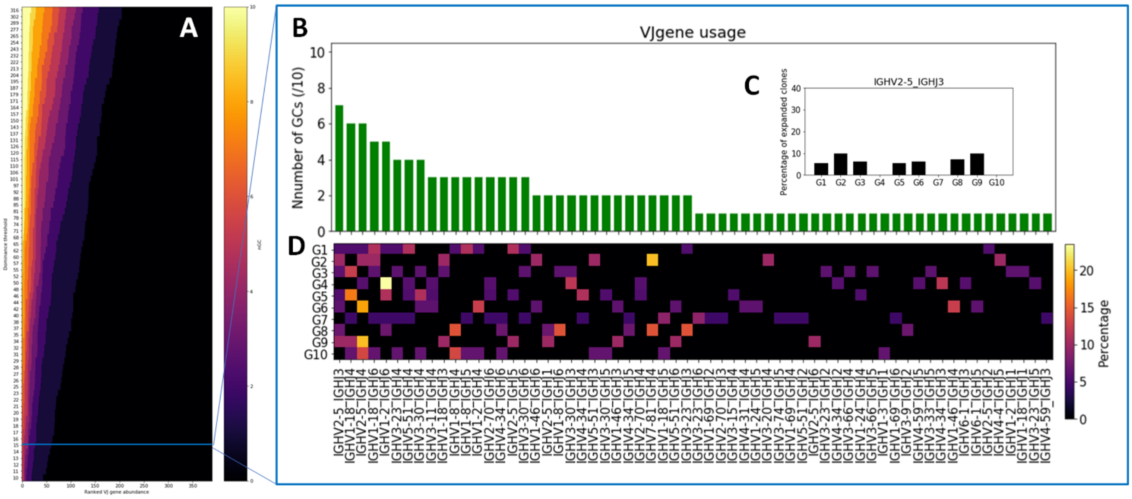

Figure S3: VJ gene usage combination heatmap. (A) VJ heatmap computed from the observed germinal center, where the $x$ axis represent the *dominance* threshold used to define the expanded clones and the $y$ axis is the histogram of the number of GCs each VJ gene is found in. (B) Histogram depicting the number of GCs for which a given V and J gene combination is found within the 15 most abundant clones of that GC, ranked from most to least abundant. (C) Histogram showcasing the usage of the most abundant VJ gene combination by the 15 most abundant clones in each GC. (D) Heatmap where one pixel represents the percentage of a given V gene being among a given GC's 15 most abundant clones.

### 4 The most abundant clones and the reoccurring clones in different GCs exhibit common features

We identified the most abundant clonotypes based on identical CDR3 AA sequences, after filtering for productive sequences, functional V gene and CDR3 with no special character, through using the TRIP-tool in an R-Shinny environment [5]. In parallel, we used the Hierarchical Agglomerative Clustering (HAC) algorithm to cluster the junctional sequences. This approach relies on identical V and J gene segments and identical CDR3 lengths, as well as more than 84 % junction nucleotide sequence overlap. We identify the dominant clones that are the most abundant by the two aforementioned algorithms, and investigate their common features, such as V D and J gene usage and CDR3 length (Table 1). We observed that the most abundant clones were identified in both replicates of the same sample, supporting the notion that our methodology is reliable in identifying the expanded clones and their features (Table 2). We also identified the clones that are shared between the different GCs and we illustrate the clones that are found by both algorithms and definitions and they share common V and J genes (Table 3).

**Table 1**

| sample | CDR3 aa sequence | CDR3 length | V gene | D gene | J gene |
|---|---|---|---|---|---|
| GC4 | CARSEGITATSFDYW | 13 | IGHV2-5 | IGHD2-15 | IGHJ4 |
| GC8 | CARGVHYDYVWGSFHWYGLDVW | 20 | IGHV1-2 | IGHD3-16 | IGHJ6 |
| GC3 | CARSNITMIGINDLW | 13 | IGHV1-18 | IGHD3-10 | IGHJ4 |
| GC5 | CAREEWLCFDLW | 10 | IGHV1-18 | IGHD3-3 | IGHJ4 |
| GC10 | CARVNNSDPGWFDPW | 13 | IGHV1-18 | IGHD3-3 | IGHJ5 |

**Table 2**

| Sample | GC | Major clonotype | Frequency (%) |
|---|---|---|---|
| 76 | GC1 | CARDPLGHWYDDSGTGGYW | 7.50 |
| 77 | GC1 | CARDPLGHWYDDSGTGGYW | 6.28 |
| 82 | GC3 | CARSNITMIGINDLW | 14.80 |
| 83 | GC3 | CARSNITMIGINDLW | 14.17 |
| 86 | GC4 | CARSEGITATSFDYW | 21.73 |
| 87 | GC4 | CARSEGITATSFDYW | 12.23 |
| 88 | GC5 | CAREEWLCFDLW | 13.80 |
| 89 | GC5 | CAREEWLCFDLW | 15.13 |
| 94 | GC8 | CARGVHYDYVWGSFHWYGLDVW | 20.92 |
| 95 | GC8 | CARGVHYDYVWGSFHWYGLDVW | 15.51 |
| 98 | GC10 | CARVNNSDPGWFDPW | 29.59 |
| 99 | GC10 | CARVNNSDPGWFDPW | 33.02 |

**Table 3**

| V-gene | Reoccurring clonotypes CDR3 AA seq | J-gene | GCs found | Frequency |
|---|---|---|---|---|
| IGHV1-18 | AREGIVETSMYKYYYGMDV | IGHJ6 | GC4<br>GC5<br>GC10 | 0,138<br>0.198<br>0.003 |
| IGHV1-18 | VRHEALSDYREIFDT | IGHJ3 | GC1<br>GC2 | 0.007<br>14.126 |
| IGHV1-18 | ARDVGGDTSWYNPGPGMDV | IGHJ6 | GC1<br>GC2<br>GC8 | 0.495<br>0.004<br>0.47 |
| IGHV2-5 | ARSEGITATSFDY | IGHJ4 | GC4<br>GC5 | 27.5556<br>0.0137 |
| IGHV2-5 | ARWGEYQDAFDI | IGHJ3 | GC1<br>GC2 | 0.0082<br>16.8594 |

Figure S4: Common features of the most abundant (Table 1) and shared clones (Table 3) in different GCs. In Table 3 the frequency corresponds to the average frequency of the replicates. The highlighted cells represent the common features of the most abundant clones and the clones identified in different germinal centers.

### 5 Identifying common binders

We optimize the *CDRs similarity*, *Paratype* [6] and *Ab-Ligity* [7] framework, described in the methods section of the main text, with a Pertussis toxin (PTx) binders dataset [6]. The dataset contains 1290 antibodies sequences from mice immunized with PTx. Sequences were annotated as PTx-binding or PTx-non-binding using using homogeneous time-resolved fluorescence (HTRF) and surface plasmon resonance (SPR). As there are multiple epitopes on the PTx antigen, most binder pairs will not be predicted to bind the same epitope. Thus, each method's performance is defined as follow:

- True positive ($TP$): A PTx-binding sequence that was identified by another PTx-binding sequence.
- False positive ($FP$): A non-PTx-binding sequence that was identified by a PTx-binding sequence.
- False negative ($FN$): A PTx-binding sequence that was not identified by any PTx-binding sequence.
- True negative ($TN$): Cannot be evaluated as only information about PTx is known, but there could be other antigens.

Prediction performances for different similarity threshold are evaluated in term of $Precision = \frac{TP}{TP+FP}$ and $Recall = \frac{TP}{TP+FN}$. The optimal threshold of each framework is taken as the one that maximizes the $Fscore = 2\ \frac{Precision \times Recall}{Precision + Recall}$ (Figure S5).

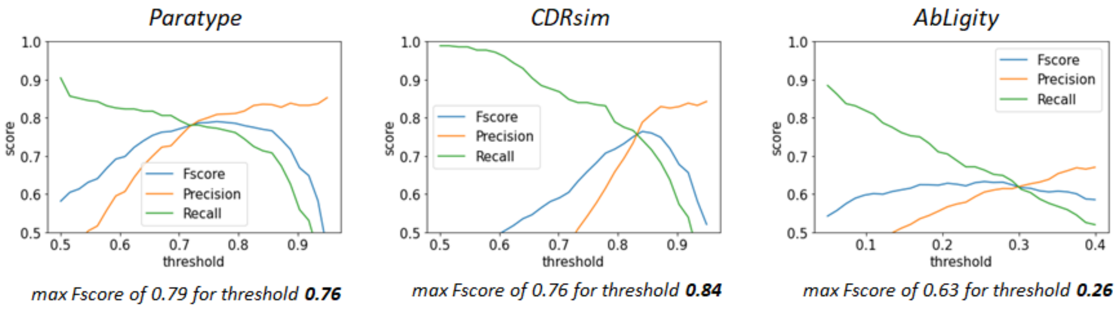

Figure S5: Threshold optimization for the *Paratype*, *CDRsim* and *AbLigity* similarity frameworks on the PTx binder dataset. The obtained thresholds for maximizing the Fscore are shown in bold.

As most of the predicted binders are *"easy"* due to being from the same clone, we further assess the prediction performances by excluding predictions of binding pairs coming from the same clone, i.e. if they share the same V gene, same J gene, same CDRH3 length and more than 84% sequence identity in the CDRH3. After removing these pairs, the accuracy of these three frameworks is significantly worse (Table S1). To improve the precision of our prediction, we combine these methods and we predict that two antibody sequences will bind to the same epitope if the similarity is above the threshold for two of the three similarity metrics. As shown in Table S1, this approach significantly improves the precision to more than 70%, but is unable to identify most of the binders, emphasized by a recall below 10%.

| | CDRsim (C) | Paratype (P) | AbLigity (A) | C and P | C and A | P and A |
|---|---|---|---|---|---|---|
| Precision | 0.314 (11/35) | 0.407 (11/27) | 0.134 (16/119) | 1.00 (8/8) | 0.70 (7/10) | 0.75 (6/8) |
| Recall | 0.129 (11/85) | 0.129 (11/85) | 0.188 (16/85) | 0.094 (8/85) | 0.082 (7/85) | 0.071 (6/85) |
| Fscore | 0.183 | 0.196 | 0.157 | 0.172 | 0.147 | 0.129 |

Table S1: CDRsim, Paratype and AbLigity performances after excluding antibody pairs from the same clone. Combining the frameworks yields higher precision. The predictions are performed with the thresholds obtained after optimization, i.e. 0.84 for CDRsim, 0.76 for Paratype, and 0.26 for AbLigity.

#### 6 Dominant clones paratope similarity across vs within GCs

As a way to study the paratope similarity of dominant clones within an individual GC versus across different GCs, we defined the paratope distance between clone pairs as (paratype distance + CDRs distance)/2. To avoid biases due to the same clone being found across different GCs, we did not consider clone pairs with the same V gene and J gene. As shown in the Figure S6, while the distances seem relatively similar to each other, a difference can be observed, with a 30% higher occurrence in the low paratope distance part for clones in the same GC compared to different GCs.

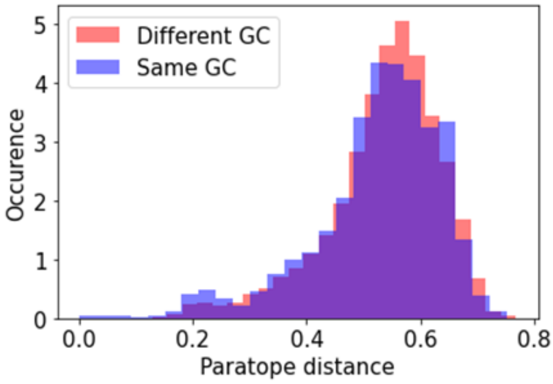

Figure S6: Distributions of paratope distances for all dominant clones pairs in the LN in the same GC (blue) and in different GC (red), with a ranking threshold of 100 in their respective GC.

#### 7 Epitope convergence model

Thanks to the framework described in the previous section, we are able to predict whether two given clones evolve toward binding the same epitope with good precision. With the same procedure used for estimating species richness in a biosystem as described in the main text, we estimate the number of epitope by considering a dominant clone as an observation and epitopes as species. First, we cluster all clones within functionality groups, such that each clone in a given group will be predicted to bind the same epitope as at least one other clone of the same group. Then, an "epitope sample" is computed for each Germinal center, where the number of clones binding each epitope is tracked. In Figure S7, we depict the epitope rarefaction curve of the 10 GCs, and the distribution of the number of clones binding each epitope. As most of the epitopes are only associated to one clone, the epitope richness in GCs is likely to be quite high. Applying the Chao [8,9] formula to the obtained sample yields.

$$N_{\text{epitopes}} = N_{\text{clusters}} + \frac{f_1\,(f_1 - 1)}{2\,(f_2 + 1)}, \tag{3}$$

where $f_1$ is the number of clusters containing exactly one clone, and $f_2$ the number of clusters containing exactly two clones. By taking the median values across GCs, we obtain an estimation of $\sim 1000$ epitopes per GC.

With a similar approach, we also estimate the number of epitopes present in the entire lymph node by looking at shared targeted epitopes across GCs. For each epitope, we compute the number of GCs in which at least one clone binding that epitope is detected. The obtained distribution is shown in Figure S8. As obtained for GCs, the number of epitopes present in the Lymph node is likely high because the majority of epitopes are found in only one GC. Applying the Chao formula gives an estimate of $\sim 5000$ epitopes present in the lymph node.

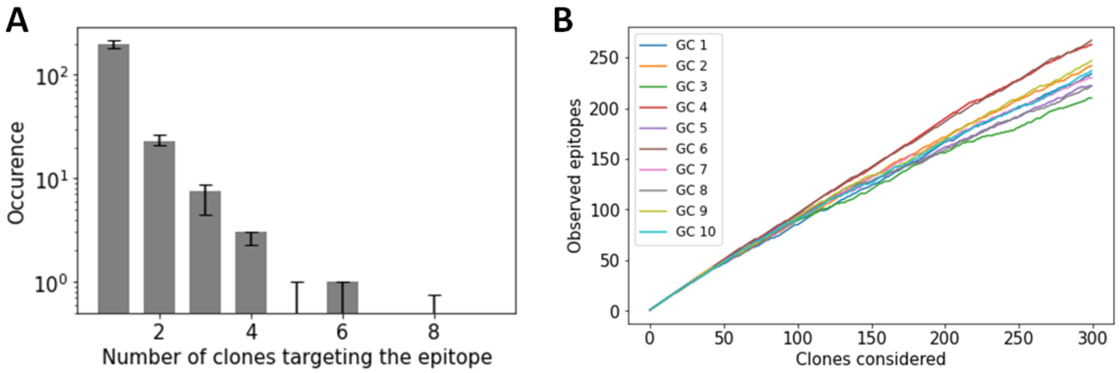

Figure S7: Epitope diversity in germinal centers where the 500 most dominant clones in each GC are considered. (A) Number of epitopes with $N$ clones in their cluster. The bar plots represent the median across GCs and the error bar corresponds to the first and third quartile. (B) Epitope accumulation curve of each GC.

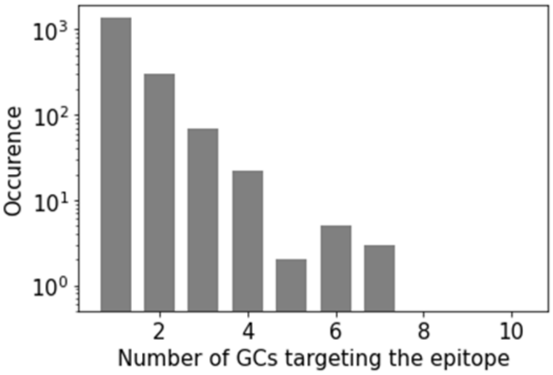

Figure S8: Distribution of the number of GCs in which a clone reacting to each epitope are found.

### 8 Supplementary figures

A Before LCM

B After LCM

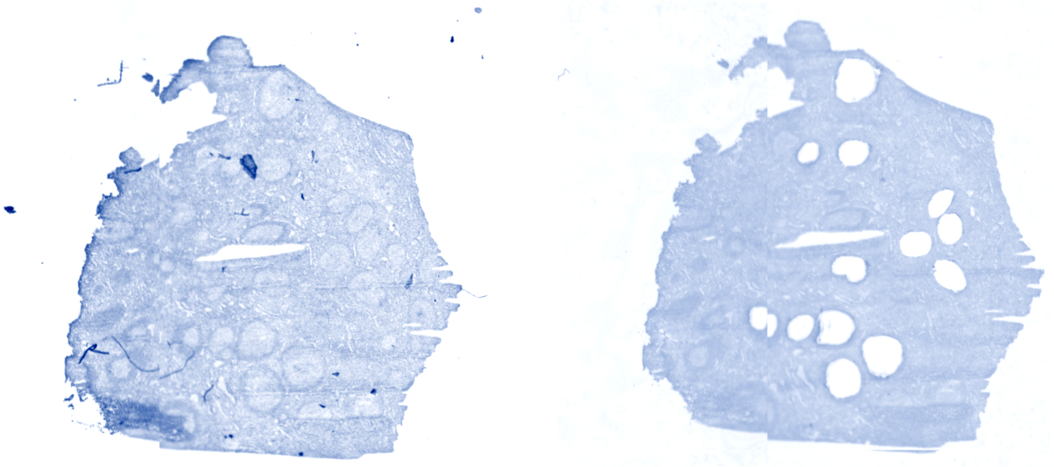

C

| | Nested-Multiplex PCR | |
|---|---|---|
| **Primer** | **Sequence** | **ROUND** |
| VH1_LE | ACCACCATGGACTGGACCTGGAGG | 1 |
| VH1B_LE | ACCACCATGGACTGGACCTGGAGM | 1 |
| VH2A_LE | ACCACCATGGACACACTTTGCTMCAC | 1 |
| VH2B_LE | ACCACCATGGACATACTTTGTTCCAAC | 1 |
| VH3_LE | ACCACCATGGAGTTTGGGCTGAGC | 1 |
| VH4_LE | ACCACCATGAAACACCTGTGGTTCTT | 1 |
| VH5_LE | ACCACCATGGGGTCAACCGCCATC | 1 |
| VH6_LE | ACCACCATGTCTGTCTCCTTCCTC | 1 |
| JH6_constant | CTTACCTGAGGAGACGGTGACC | 1 |
| VH1-FR1 | TCGTCGGCAGCGTCAGATGTGTATAAGAGACAGGGCCTCAGTGAAGGTCTCCTGCAAG | 2 |
| VH2-FR1 | TCGTCGGCAGCGTCAGATGTGTATAAGAGACAGGTCTGGTCCTACGCTGGTGAAACCC | 2 |
| VH3-FR1 | TCGTCGGCAGCGTCAGATGTGTATAAGAGACAGCTGGGGGTCCCTGAGACTCTCCTG | 2 |
| VH4-FR1 | TCGTCGGCAGCGTCAGATGTGTATAAGAGACAGCTTCGGAGACCCTGTCCCTCACCTG | 2 |
| VH5--FR1 | TCGTCGGCAGCGTCAGATGTGTATAAGAGACAGCGGGGAGTCTCTGAAGATCTCCTGT | 2 |
| VH6-FR1 | TCGTCGGCAGCGTCAGATGTGTATAAGAGACAGTCGCAGACCCTCTCACTCACCTGTG | 2 |
| JH6_constant | GTCTCGTGGGCTCGGAGATGTGTATAAGAGACAGCTTACCTGAGGAGACGGTGACC | 2 |
| **NGS-Miseq-Adaptors** | | |
| Forward overhang: | TCGTCGGCAGCGTCAGATGTGTATAAGAGACAG | |
| Reverse overhang | GTCTCGTGGGCTCGGAGATGTGTATAAGAGACAG | |

Figure S9: Extensive Experimental Method, (A) The Human LN tissue used on a PEN slide before and after LCM is shown. The fast tissue specimen overview was automatically created by the Leica PALM-LMD6 with 1.25X magnification. The scale, colour and the brightness have been optimized for better clarity (B) Primer set pools (10 uM per primer) were prepared for the LD and the FR1 region PCR amplification for round 1 and round 2, respectively

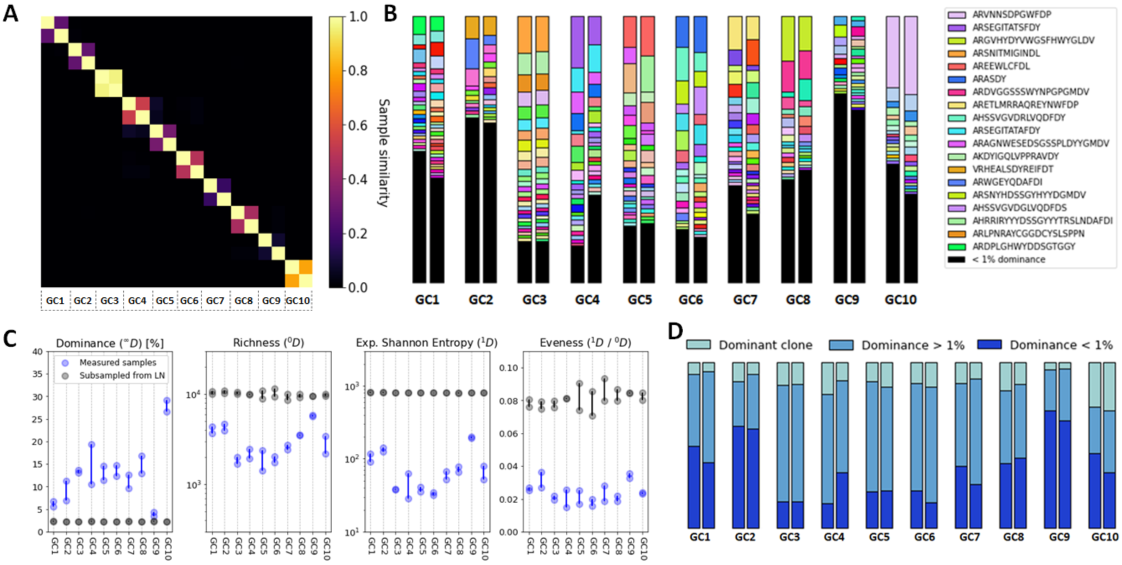

Figure S10: Samples diversity analysis with CDR3. (A) Sørensen–Dice similarity between samples in term of CDR3 abundance. (B) CDR3 abundance across samples, only the 20 most abundant CDR3s across samples are shown in the legend. (C) Diversity analysis across samples in term of dominance, richness, Shannon entropy and evenness, where ${}^{q}D$ corresponds to the Hill's unified notation. To highlight the relevance of studying GCs individually, each sample (blue) was compared to an artificial sample of equivalent size sub-sampled from sequences in the LN (grey). (D) Proportion of sequences belonging to the dominant clone, expanded clones and non-expanded clones in each sample.

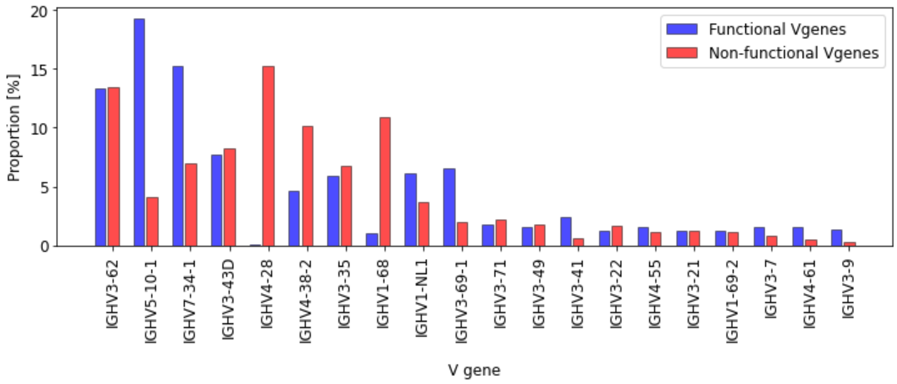

Figure S11: V gene frequency in functional and non-functional clones.

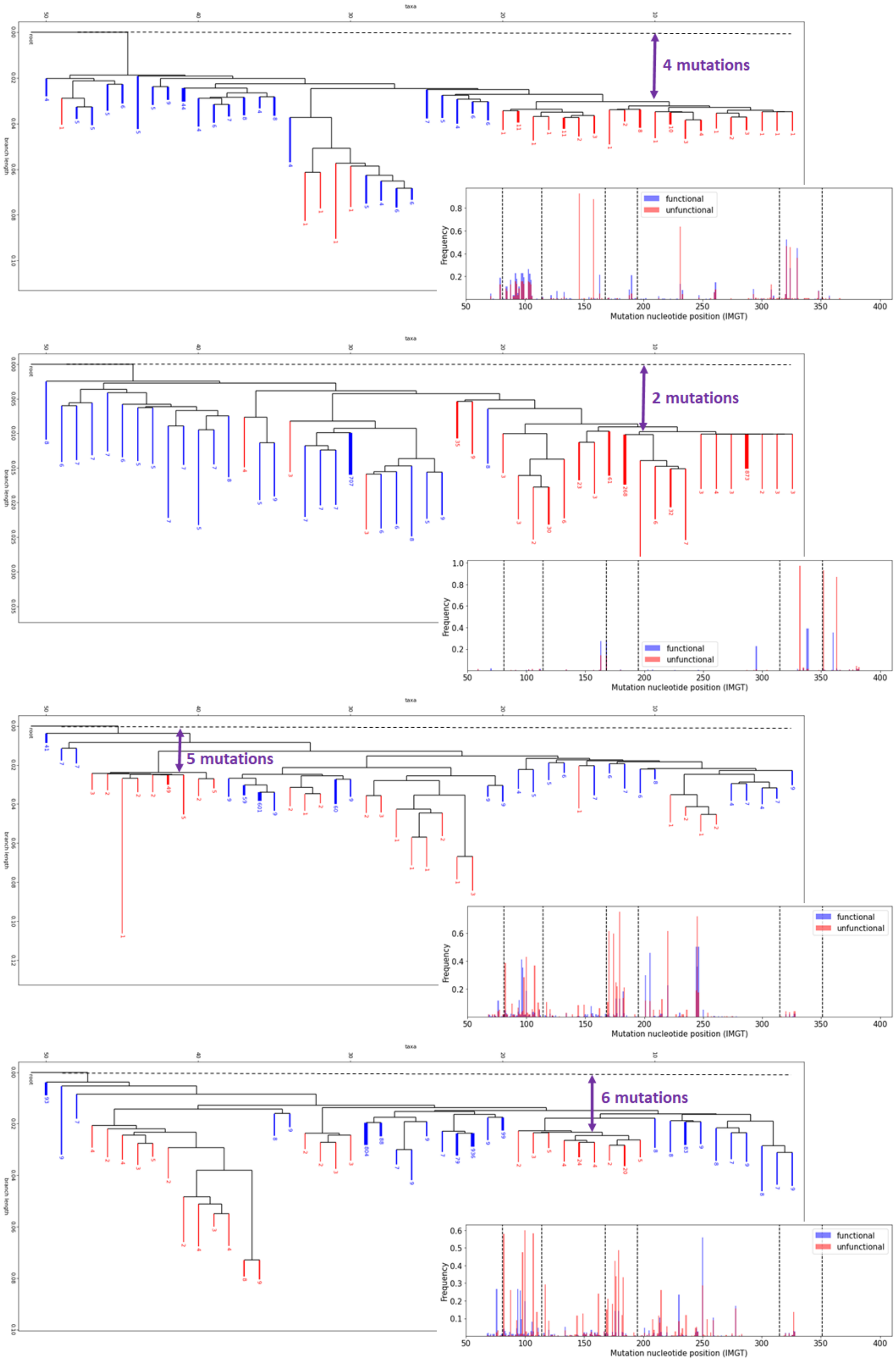

Figure S12: Phylogeny reconstruction of some representative F&NF clones.

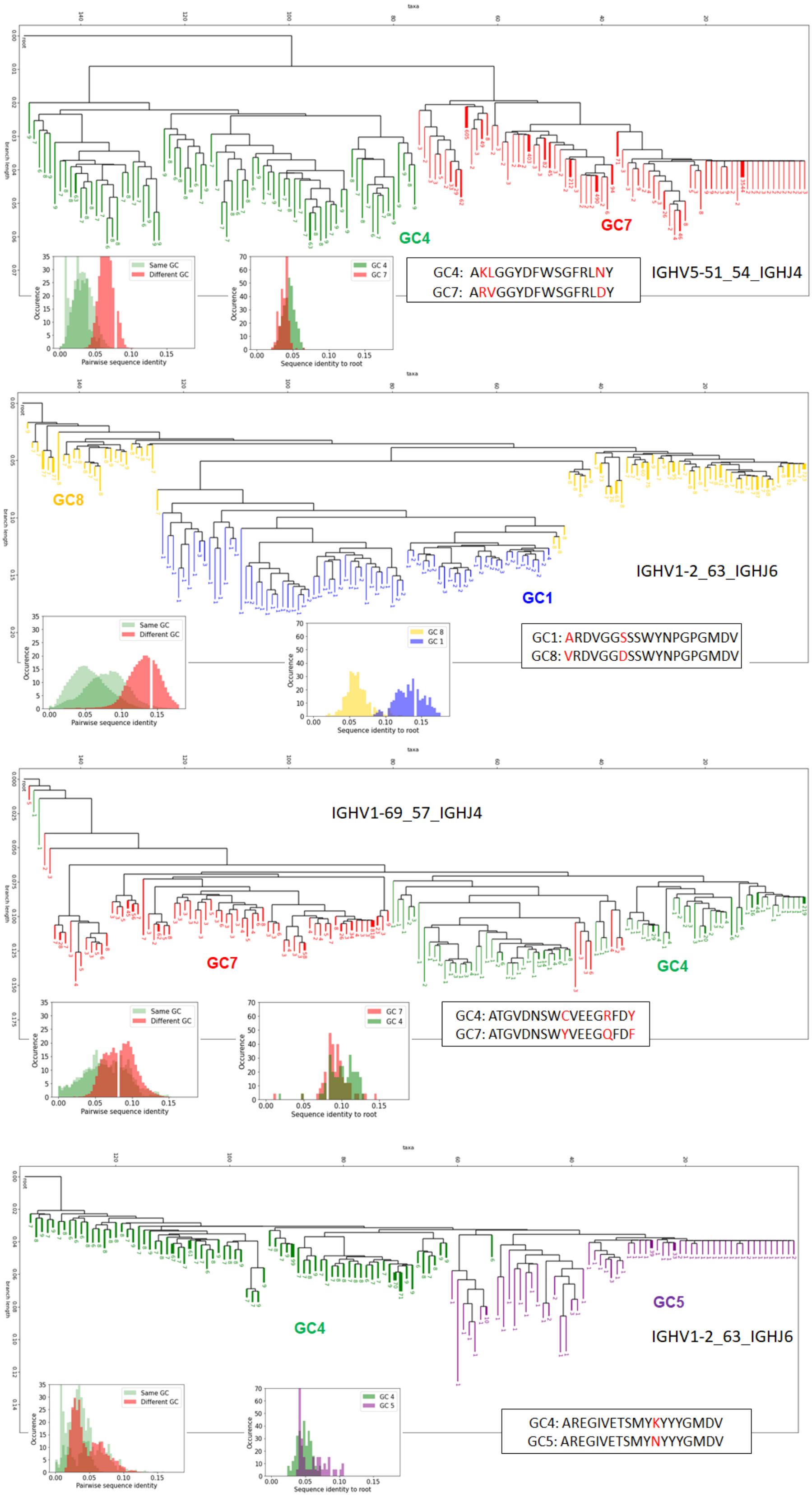

Figure S13: Phylogeny reconstruction of some representative shared clones across GCs. Only the 300 most abundant branches are shown for visual clarity
